# Multiplexed sequence-resolved screening of transient DNA hybridization for programmable nanotechnology

**DOI:** 10.64898/2026.08.25.746935

**Authors:** Carolien Bastiaanssen, Ran Huo, Patrick Irmisch, Archana Sivaraman, Ralf Seidel, Kristin S. Grußmayer, Chirlmin Joo

## Abstract

DNA-based technologies rely on short, transient hybridization events, but selecting sequences with desired kinetic properties remains largely empirical because hybridization kinetics are difficult to predict from sequence and slow to measure one sequence at a time. Here, we introduce SPARXS-Hyb, an implementation of SPARXS (Single-molecule Parallel Analysis for Rapid eXploration of Sequence space) for multiplexed sequence-resolved screening of DNA hybridization. Using a surface-immobilized docking-strand library and a quencher-labelled imager-strand library, we screened 128 different DNA sequences in a single kinetic measurement, exposing all sequences to identical experimental conditions. This multiplexed approach removes a major confounding factor of serial measurements, allowing sequence-dependent differences to be compared directly. The resulting dataset reveals sequence-dependent transient binding behaviours and enabled us to identify a sequence with which an order-of-magnitude higher sampling rate can be achieved in DNA-PAINT (DNA points accumulation for imaging in nanoscale topography), a super-resolution microscopy technique based on DNA hybridization. By enabling multiplexed screening across a sequence library, SPARXS-Hyb provides a route to kinetics-guided sequence selection for programmable transient interactions in DNA nanotechnology.

## Introduction

Core biomolecular techniques such as polymerase chain reaction and southern blotting rely on the process of DNA hybridization. In recent years, creative use of the programmability of DNA hybridization has led to major advances in nanoscale technologies^1^. DNA origami enables the assembly of custom-shaped nanoscale structures for nanophotonics, DNA computing, and targeted drug delivery^2,3^. Beyond structural nanotechnology, DNA hybridization is exploited in advanced imaging methods. An example is the super-resolution microscopy technique DNA-PAINT (DNA points accumulation for imaging in nanoscale topography)^4^. It makes use of fluorescently labeled short DNA oligonucleotides (imager strands) that transiently hybridize to complementary oligonucleotides (docking strands) which are attached to target molecules. DNA-PAINT super-resolution imaging requires long acquisition times which makes it slow compared to other super-resolution microscopy techniques. The hybridization frequency can be increased by using higher imager concentrations; however, this increases the background signal. Recently, the introduction of repetitive docking strands has improved image acquisition speed^5–7^. These docking strands present multiple binding sites for the imager strand per target and thereby increase the hybridization frequency. However, for sub-10-nm scales, repetitive docking strands induce spatial blurring due to a combination of factors related to the increased length and the presence of multiple binding sites^8^. Another strategy to accelerate DNA-PAINT imaging is by optimizing the imager sequence^6^, as both the hybridization rate and the stability of the duplex depend largely on the DNA sequence.

This sequence dependence raises a central design challenge: while DNA hybridization thermodynamics can be predicted with considerable accuracy^9,10^, sequence-dependent hybridization kinetics remain much harder to forecast. Recently, several groups have begun tackling the harder problem of predicting hybridization kinetics. Models from Hata et al., Zhang et al., and Hertel et al. have provided valuable insights, yet accurately forecasting kinetics, especially for short, transiently binding sequences remains challenging^11–13^.

As a result, candidate sequences are typically selected by a combination of predicted thermodynamic stability and empirical design rules and are then validated individually. This has led to one of the fastest sequences up to date for DNA-PAINT super-resolution imaging^6^. However, this approach explores only a small fraction of the available sequence space. Moreover, because candidates are tested in separate measurements, apparent kinetic differences can be confounded by variations in surface preparation and imaging conditions. Two approaches developed in parallel, SPARXS (Single-molecule Parallel Analysis for Rapid eXploration of Sequence space)^14,15^ and MUSCLE (multiplexed single-molecule characterization at the library scale)^16,17^, overcome this limitation by coupling single-molecule fluorescence microscopy with next-generation sequencing, yielding sequence-resolved single-molecule fluorescence time traces. To enable direct selection of optimal hybridization sequences for nanotechnology applications, we developed a single-molecule multiplexed DNA hybridization assay based on SPARXS termed SPARXS-Hyb (SPARXS for DNA Hybridization). In SPARXS-Hyb, different DNA sequences are measured in the same single-molecule experiment and then identified by sequencing, allowing their hybridization behaviour to be compared under identical experimental conditions. We kinetically screened a library of 128 DNA sequences and identified a sequence (SH6) that enables DNA-PAINT super-resolution imaging with an order of magnitude higher sampling rate than the current standard in the DNA-PAINT field (PS3)^6^ and validated it using cellular DNA-PAINT super-resolution imaging.

### Multiplexed sequence-resolved DNA hybridization assay

In a SPARXS-hyb experiment, a library of DNA sequences is immobilized on the surface of a next-generation sequencing flow cell, a complementary library is added in solution, followed by single-molecule fluorescence microscopy imaging, and subsequent identification of each molecule by sequencing (Fig. 1a). We focused on 7-nucleotide sequences because they are short enough to support rapid dissociation and repeated binding cycles, but sufficiently stable to generate detectable single-molecule events under DNA-PAINT-like conditions. This length is also directly relevant to DNA-PAINT, as PS3, a widely used speed-optimized imager sequence, is 7 nucleotides long, and previous rational screening of 6-, 7- and 8-mers identified 7-mers among the best-performing candidates^6^. Since secondary structures negatively impact DNA-PAINT performance by slowing hybridization kinetics and reducing binding event frequency^18^, we decided to focus the library on all possible 7-nucleotide combinations of adenine and guanine on the surface (docking library) and of cytosine and thymine in solution (imager library). This restricted 7-mer library provides a controlled demonstration of sequence-resolved screening while retaining a design space that is directly relevant to transient DNA-PAINT interactions.

**Fig. 1.**
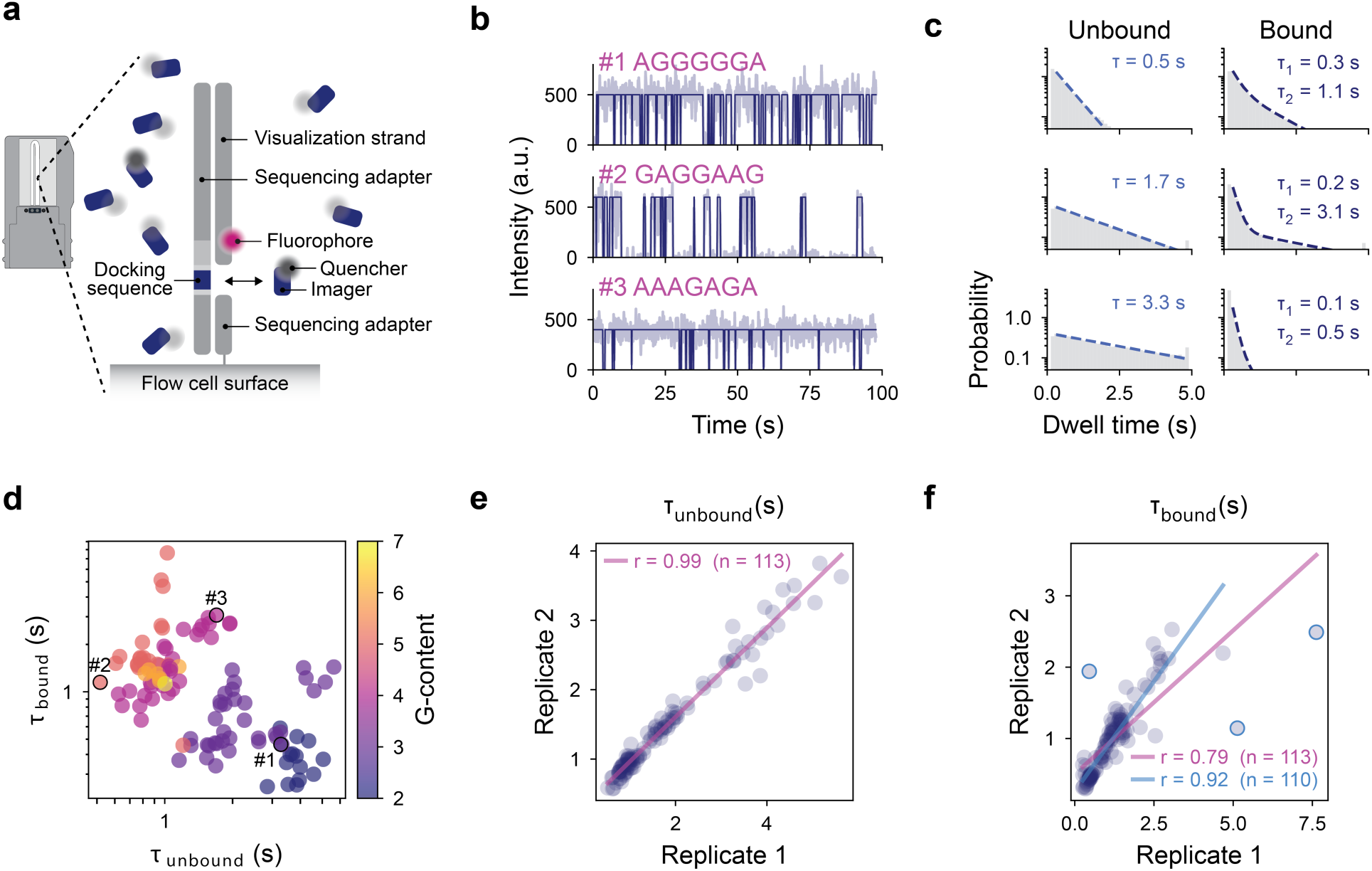
Parallel kinetic profiling of 128 DNA sequences with SPARXS-hyb. **a,** Schematic of the SPARXS-hyb sample in the sequencing flow cell. **b,** Sequence-coupled fluorescence time traces obtained by SPARXS-hyb. Representative traces are shown for three different sequences with the raw data in light and the hidden Markov model classification in dark. Hybridization events are observed as a sharp decrease in intensity. **c,** Probability density distribution of the observed bound and unbound dwell times, τ_bound_ and τ_unbound_, for the three sequences in b with fit. The number of dwell times on which the distributions are based are reported in Table S1. **d,** Scatter plot of τ_bound_ versus τ_unbound_. The markers are colour coded for the number of guanines in the docking sequence and the highlighted sequences correspond to the sequences in b and c. **e,** Scatter plot comparing τ_unbound_ between duplicate SPARXS-Hyb experiments. **f**, Scatter plot comparing τ_bound_ between duplicate SPARXS-Hyb experiments. A second correlation is shown in blue where the three sequences with the largest relative 95% confidence interval were excluded. For d, e and f, sequences with an event frequency below 0.05 Hz were excluded, giving a total of 113 sequences.

The immobilized docking library was visualized with a stably hybridized common visualization strand containing a fluorophore (Cy3). For each imager sequence we used a concentration of 10 nM to obtain enough binding events within the single-molecule observation window of 80 seconds. Because the 128-member imager library therefore exceeded 1 µM in total concentration, direct fluorescent labeling of all imagers would generate prohibitively high background for single-molecule TIRF imaging. We therefore labeled the library with a non-fluorescent quencher (BHQ2). In this design, an unbound docking strand gives a high signal, and hybridization of an imager from solution causes a sharp decrease in the intensity (Fig. 1b). After automated imaging over the entire flow cell surface with a total imaging time of approximately 91 hours, the immobilized library is sequenced. The fluorescence and sequencing datasets are coupled, enabling us to analyse the sequence dependence of the hybridization kinetics of the library.

### Multiplexed kinetic profiling of 128 DNA sequences

In a single SPARXS-Hyb experiment, using a MiSeq v3 chip, 4080 movies of 80 seconds were acquired, covering a total area of 15.7 mm^2^. This yielded 5.1 million single-molecule fluorescence time traces and sequencing returned 4.6 million sequences originating from the imaged surface. Coupling of these two datasets led to 1.3 million sequence-coupled fluorescence time traces with a coverage of over a thousand traces per library sequence (Fig. S1). We collected the traces per sequence and classified bound and unbound states using a hidden Markov model (see Methods), assigning the high-intensity state to the unbound docking strand and the low-intensity state to imager-bound states. From this analysis, we obtained probability density distributions for the bound and unbound dwell times (Fig. 1c, Fig. 2). Sequences with very few detected events (an event frequency below 0.05 Hz) were excluded from further analysis since below this threshold the number of events within our experiment time was too low to construct a reliable dwell time probability density distribution and resulted in unreliable fits (Fig. 2, sequences shaded in grey; Fig. S2).

**Fig. 2.**
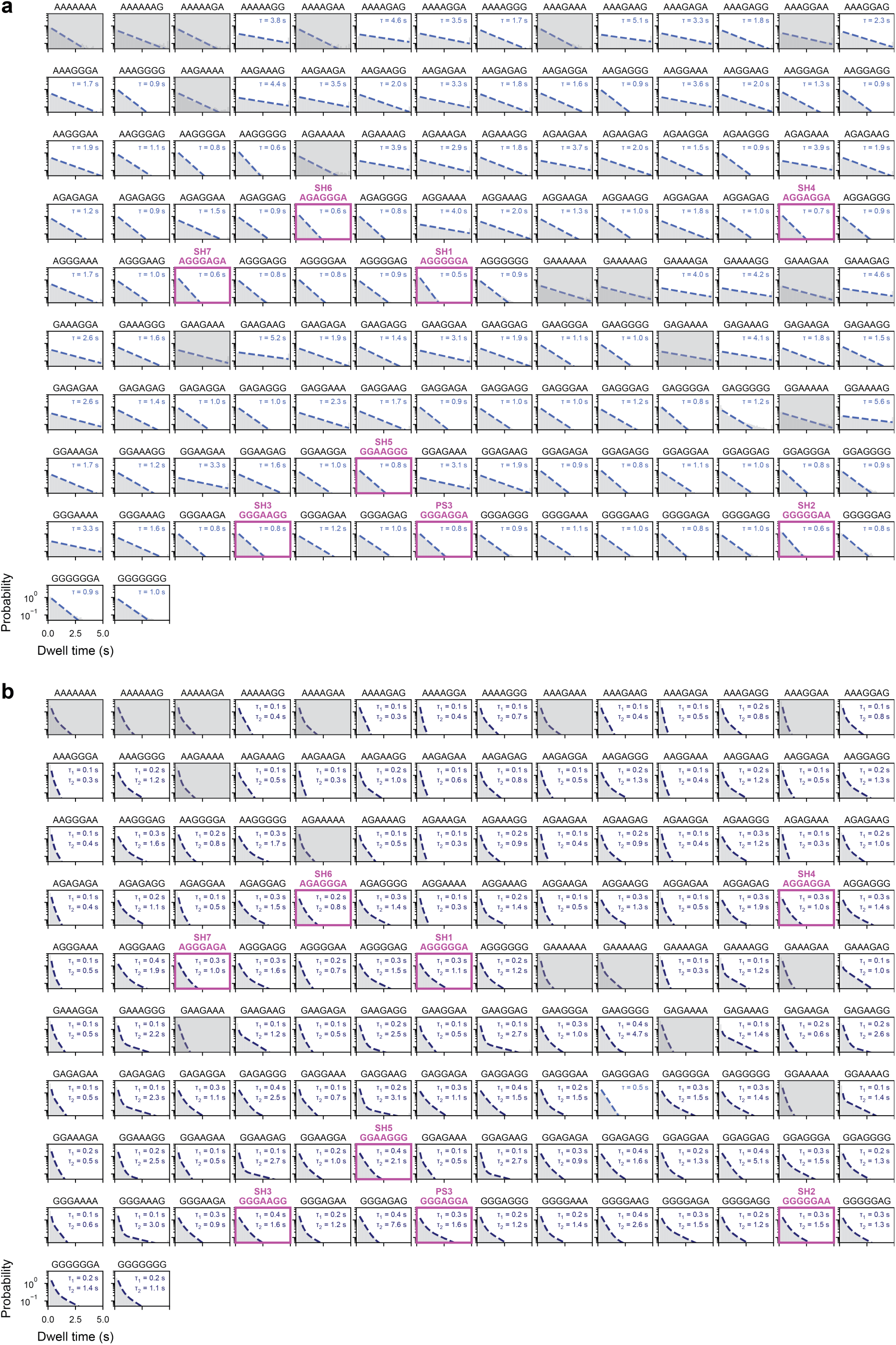
Probability density distributions of the observed unbound (a) and bound (b) dwell times. Single exponential fits are shown in light blue and double exponential fits in dark blue. Sequences with a binding frequency below 0.05 Hz are shaded in grey, and their dwell times were not used for further analysis. The SH sequences and PS3 are highlighted in magenta. The fits and the number of dwell times on which the distributions are based are reported in Table S1.

The probability density distributions of the bound dwell times revealed the intrinsic heterogeneity of the interactions. The distributions of the unbound dwell times were well-described by single exponential decays, providing an apparent association rate for each docking sequence. In contrast, the bound dwell-time distributions were more adequately described by a double-exponential decay and often exhibited a pronounced separation between short-lived and long-lived binding populations. This complexity is expected from the design of the SPARXS-Hyb assay, in which binding events can arise from both the perfectly matching imager and mismatching imagers. We therefore interpreted the dwell-time distributions as descriptive observables rather than direct measurements of sequence-specific rates. The longer component was taken as the observed bound dwell time based on the expectation that a perfectly matching imager binds longer than the mismatched imagers (Fig. S3). The resulting fits showed that our library covers a broad dynamic range of bound and unbound dwell times (Fig. 1d; Table S1), from 0.52 to 5.63 for unbound and from 0.25 to 7.63 for bound dwell times.

We also performed a replicate SPARXS-Hyb experiment, which shows a strong correlation with the first one (Fig. 1e and f). We further validated our approach by comparing the association rates of all mirror sequences (i.e. pairs of sequences where the 5’-3’ direction of one sequence equals the 3’-5’ direction of the other) in the library, which likewise showed a strong correlation for the observed association rates (Pearson correlation of 0.96, Fig. S4).

### Sequence-resolved rate analysis identifies fast transient binders

Although the mixed imager library complicates the interpretation, we aimed to relate the observed kinetics to thermodynamic properties. To this end, we used NUPACK^19^ and calculated the free-energy change upon hybridization for each imager sequence to its complement. Plotting the apparent association rate (k_on_) obtained from the unbound dwell times as a function of the predicted free-energy change (Δ*G*) provided a clear correlation across the library (Fig. 3a). Furthermore, this analysis provided an estimate for the minimum free-energy change required for us to observe binding, for which we determined the threshold to be −14.5 k_B_T, as sequences above this value did not pass the event-frequency threshold.

**Fig. 3.**
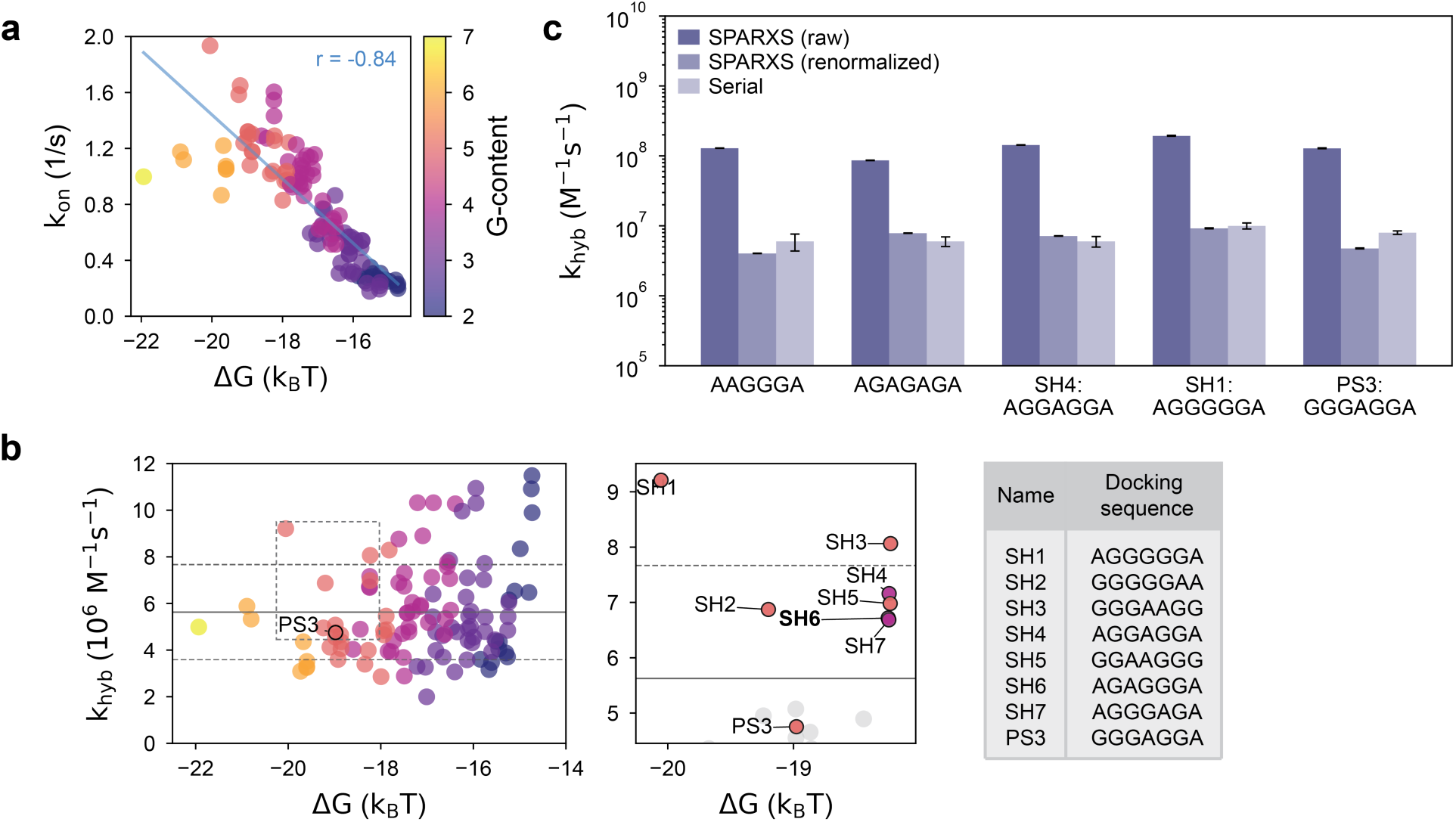
Free-energy dependence and renormalization of SPARXS-Hyb association rates. **a,** Scatter plot of the observed association rates (k_on_) versus the predicted free-energy change upon hybridization (Δ*G*). The colour of the data points indicates the number of guanosines in the docking sequence. **b,** Scatter plot of the renormalized association rates with the mean and one standard deviation indicated by the solid and dashed lines respectively. Colouring is the same as in a. The outlined area is shown as a zoom-in on the right with the candidate sequences SH1-7 and PS3 in the table to the right. Sequences with an event frequency smaller than 0.05 Hz were excluded in both a and b, leaving 113 included sequences. **c,** Bar plot comparing hybridization rates obtained through serial measurements and SPARXS. For SPARXS, both the raw and renormalized hybridization rates are shown. Serial data is based on three independent replicates with error bars indicating the standard deviation of the mean. The SPARXS data is based on replicate 1 and the error bars indicate the 95% confidence interval based on bootstrapping. Statistics of the SPARXS data can be found in Table S1.

We next used this free-energy threshold to estimate, for each docking sequence, how many imagers in the library would be sufficiently stable to contribute to the observed binding events. This number varied substantially across the library. Weak sequences have only a few mismatched binders but strong sequences with many guanines have up to 30 different imagers that can bind (Fig. S5). Considering that the observed association rate is the sum over all potential binders, and assuming that each of the potential binders for a certain docking sequence has a similar association rate, we renormalized the association rate for each docking sequence by dividing the observed rate by the total number of potential binders (Fig. 3b).

We tested our approach by comparing the renormalized rates with rates acquired through serial control experiments that contain only the matching pair of docking and imager strands (Fig. 3c). The agreement between the multiplexed and serial measurements supports the use of the renormalized rate as a practical predictor of matched-pair association behaviour. Additionally, the renormalized association rates scatter around a common value of approximately 6 · 10^6^ *Ms*^−1^ which is consistent with values previously reported^12,13^. Interestingly, there are measurable differences in association rates between sequences within similar predicted free energies, indicating that SPARXS-Hyb can select fast binders beyond what would be expected from thermodynamic stability alone.

### Super-resolution imaging with an order-of-magnitude higher sampling rate

We subsequently used the SPARXS-Hyb dataset to select short, non-repetitive candidate sequences for faster DNA-PAINT imaging, potentially outperforming the current standard in the DNA-PAINT field (PS3)^6^. In search of a sequence that enables faster DNA-PAINT super-resolution imaging, we applied the following selection criteria: (1) a higher association rate than PS3 to increase the sampling rate and (2) a free-energy change upon hybridization comparable to that of PS3, to retain transient binding and avoid excessively long-lived binding events. In total, seven sequences passed our selection criteria and we named them SPARXS-Hyb-1 (SH1) to SH7. The recovery of SH4, previously identified as PS4 by rational DNA-PAINT sequence design^6^, independently supports SPARXS-Hyb as an effective approach for kinetic sequence selection.

We first tested SH1, the highest-ranked candidate; however, SH1, did not show frequent binding in cells. Because SH1 contains a stretch of four consecutive guanines, we considered that alternative structures, such as G-quadruplex formation, might compromise its cellular DNA-PAINT performance. We therefore deprioritized SH2, which also contained a G-stretch. SH4, identical to PS4, has previously been shown to be outperformed by PS3^6^. Among the remaining candidates, SH6 and SH7, which are mirror sequences, had the shortest bound dwell time combined with a high renormalized association rate, PS3-like predicted stability and no extended G-stretch. We therefore selected SH6 for cellular DNA-PAINT validation.

To test the applicability of SH6 for super-resolution imaging using DNA-PAINT, we used it to image microtubules in cells. We imaged a complete cell with SH6 using 200 pM of imager and a camera exposure time of 50 ms (Fig. 4a). Next, we compared the performance of SH6 with that of the speed-optimized sequence PS3 and found that SH6 exhibits an approximately 10-fold higher sampling rate than PS3 when reconstructing microtubule filaments. At a low imager concentration of 100 pM, SH6 exhibits blinking kinetics comparable to those of PS3 at 1 nM, and complete microtubule filaments appear after 250 seconds of acquisition time (Fig. 4b; Movie S1). When increasing the SH6 imager concentration to 250 pM, the density of emitters in each frame increased, resulting in rapid reconstruction of visible filaments after 20 seconds of acquisition, as is also demonstrated by the increased number of localizations identifiable per frame per unit area in areas of similar filament density (Fig. 4c). The lower imager concentration required for SH6 to achieve the sparse blinking conditions necessary for single-molecule localization microscopy (SMLM) has as an advantage that the background noise is reduced, leading to a higher signal-to-background ratio (SBR) thus an improved localization precision (Fig. S6, Table S2).

**Fig. 4.**
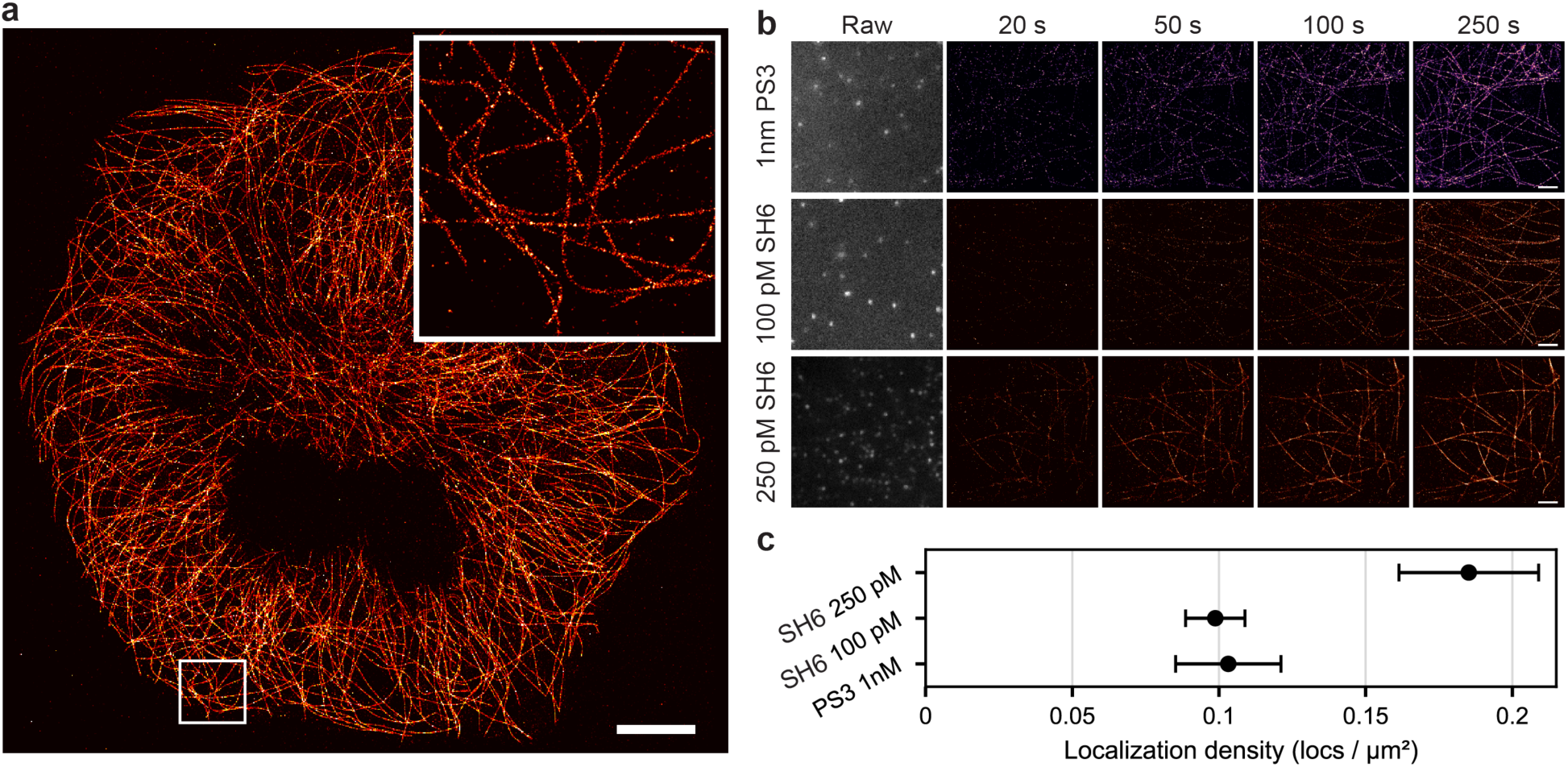
Cellular DNA-PAINT super-resolution imaging with a novel fast-sampling sequence identified by SPARXS-Hyb. **a**, Whole-cell SMLM reconstruction of microtubules imaged with 200 pM SH6. A localization precision estimation of 7.82 nm was achieved. The insert shows a zoom-in of the highlighted area and the scale bar equals 10 µm. **b**, DNA-PAINT imaging of microtubules with the sequence as indicated on the left. From left to right: one raw image frame showing emitters in the fluorescence on-state, SMLM reconstruction after 20, 50, 100, and 250 seconds of image acquisition. Scale bars equal 2 µm. **c**, Comparison of average localization density per frame between 1 nM PS3, 100 pM SH6 and 250 pM SH6. The error bars indicate standard deviation.

### Designing transient hybridization kinetics for DNA nanotechnology

The SPARXS DNA hybridization assay presented here provides a multiplexed quantitative approach for selecting short DNA sequences with desired transient hybridization behaviour. We demonstrate direct comparison of sequence-dependent binding behaviour under identical surface, buffer, and imaging conditions, with a library of 128 sequences. This is distinct from conventional serial measurements, in which apparent kinetic differences can be confounded by experiment-to-experiment variation. From the 128 sequences screened in a single experiment, we identified SH6, a sequence that outperforms PS3, the current DNA-PAINT standard. With SH6, super-resolution images can be obtained at a tenfold higher sampling rate compared to PS3, enabling the use of lower imager concentrations without sacrificing acquisition speed. This improvement directly translates to lower background noise and higher localization precision in the reconstructed super-resolution images. The systematic kinetic screening enabled by SPARXS-Hyb thus provides the means to uncover sequence variants with improved performance.

The ability to select sequences by experimentally measured transient binding behaviour is relevant beyond DNA-PAINT. Many DNA-based nanotechnologies rely on transient hybridization events whose performance is difficult to optimize by thermodynamic prediction alone. We anticipate SPARXS-Hyb to accelerate the selection of sequences in various applications such as dynamic DNA origami structures, DNA computing, and DNA walkers for tuned lifetimes and turnover rates^3,20^. Furthermore, this assay can be used to study the effects of mismatches on DNA hybridization kinetics, a key determinant for optimization of gene-editing systems like CRISPR, and hybridization-tools for detecting single-nucleotide variants (SNVs).

The platform is not limited to the 128-sequence library and can be readily scaled to larger libraries. For the replicate experiment a smaller flow cell was used which yielded a factor 10 less traces than the original experiment (Fig. S1). Since it replicated the data very well with this smaller number of traces per sequence (Fig. 1e and f), the library size can be increased by at least an order of magnitude with the larger flow cell of the original SPARXS-Hyb experiment.

The current implementation of SPARXS-Hyb is best viewed as a sequence-screening platform rather than a complete kinetic deconvolution method. Because all imager sequences are present simultaneously, observed binding events can arise from both perfectly matched and mismatched imagers. This design makes it possible to screen many sequences in parallel at useful per-sequence concentrations, but it also complicates the interpretation of bound dwell-time distributions. Further analysis of the current dataset might yield an approach to extract the dissociation rates from the mixture, but a more sophisticated approach to obtain these rates is the use of an orthogonal sequence library^21^.

Beyond its advantages for DNA-PAINT, the rapid blinking kinetics and improved signal-to-noise ratio observed with SH6 open new possibilities for other techniques that rely on fluorescence blinking through transient hybridization events. For example, super-resolution optical fluctuation imaging (SOFI) could benefit from a sequence with these characteristics. SOFI uses higher-order statistical analysis of temporal intensity fluctuations to extract sub-diffraction spatial information^22,23^ and therefore depends critically on measuring sufficient blinking statistics. Due to its enhanced blinking kinetics, SH6 could thus boost super-resolution reconstruction by SOFI. More generally, SPARXS-Hyb could be used to select sequences with specific hybridization kinetics: long on-times for prolonged tracking with single-particle tracking (SPT), very short on-times paired with high hybridization rates for minimal fluorescence photon fluxes (MINFLUX) microscopy and multiple sequences with distinct hybridization kinetics for kinetic multiplexing^24^. By enabling multiplexed sequence-resolved screening of short transient DNA hybridization events, SPARXS-Hyb provides a practical route to designing programmable binding interactions for advanced microscopy and other dynamic DNA-based technologies.

## Methods

### SPARXS-Hyb library preparation

Synthetic DNA was purchased from Ella Biotech (Germany; Table S3). The visualization strand was labeled with Cy3 Mono NHS ester (Sigma-Aldrich). For the labeling reaction, 5 μl of 200 μM DNA, 1 μl of freshly prepared 0.5 M sodium bicarbonate and 1 μl of 20 mM dye in DMSO were mixed and incubated for 6 hours at room temperature in the dark. Ethanol precipitation was performed and the labeling efficiency was determined using a spectrophotometer (DeNovix DS-11+). The labeling efficiency was approximately 100%. The final samples were obtained by hybridizing the docking and visualization strand, with an additional immobilization strand for the control experiments. Hybridization occurred in a 1:1:1 ratio in annealing buffer (10 mM Tris pH 8, 1 mM EDTA, 50 mM NaCl) by heating to 90 °C for 3 min and then slowly cooling with 1 °C every min to 4 °C.

### Flow cell preparation

The sequencing flow cell was prepared as described previously^14,15^. In short, an Illumina MiSeq flow cell was washed, closed with tape, and bleached for 5 h with a blue LED (Kessil PhotoReaction PR160L-456-EU, 50 W). For the first replicate a v3 Nano MiSeq flow cell was used and for the second replicate a v2 Nano MiSeq flow cell was used.

For serial experiments on custom made flow cells, quartz slides with a polyethylene glycol-passivated surface were prepared as described previously^25^. Each channel was incubated with 20 μl 0.1 mg/ml streptavidin (Sigma-Aldrich) for 30 s and unbound streptavidin was flushed out with 100 μl T50 (10 mM Tris-HCl pH 8.0, 50 mM NaCl). Next, 50 μl 50 pM nucleic acid sample was introduced into the chamber and incubated for 1 min, after which unbound sample was flushed out with 100 μl T50. Next, the flow cell was washed with 100 μl imaging buffer consisting of 50 mM Tris-HCl pH 8.0, 500 mM NaCl, 1 mM Trolox (6-Hydroxy-2,5,7,8-tetramethylchroman-2-carboxylic acid, Sigma-Aldrich), 2.5 mM protocatechuic acid (PCA; Sigma-Aldrich) and 0.155 U/µl protocatechuate-3,4-dioxygenase (PCD; OYC).

### Experimental set-up and data acquisition for SPARXS-Hyb

Imaging buffer with 1.28 µM imager library was added to the flow cell, after which the inlet and outlet were covered with air-tight tape (Tesa, 4965 Original) to prevent evaporation. Single-molecule imaging of the SPARXS-Hyb experiments was performed on an objective-type total internal reflection fluorescence (TIRF) microscope (Nikon Eclipse Ti2). The microscope was equipped with a 100x oil immersion objective (Nikon CFI Apochromat TIRF 577 100XC Oil) through which the sample was excited and imaged. Excitation occurred in a 360-degree fashion with a 561 nm laser (Gataca iLaunch system, Gattaca iLas2). The collected signal was filtered with two filters (FF01-609/54 and FF01-600/52, Semrock), both being held in a splitting module which was used in bypass mode (OptoSplit 2). Finally, the signal was projected on a CCD camera (Andor iXon Ultra 897). The microscope was equipped with an automated stage and automated focusing system, enabling automated image acquisition using the MetaMorph software.

### Experimental set-up and data acquisition for serial DNA hybridization measurements

Imaging buffer with 10 nM of the indicated imager was added to the flow cell. Serial experiments on quartz were performed on a custom-built prism-type TIRF microscope (Nikon Eclipse Ti2). For excitation, a 550 mW 532 nm laser was used (L4Cc-CSB-1311, Oxxius). A 60x water immersion objective (CFI Plan Apochromat VC 60x WI, Nikon) was used to collect the emission signal, which was subsequently filtered using a quad-notch filter in the turret (NF03-405/488/532/635E-25, Semrock). In an external emission box, the signal was split into two channels using a dichroic mirror (T635lpxr, Chroma) and further filtered with emission filters (ET585/65m for the Cy3 and ET655LP for the Cy5 emission signal, Chroma) before being projected onto a CMOS camera (Prime BSI sCMOS, Photometrics) using a dichroic mirror (T635lpxr, Chroma). Movies were acquired using NIS-Elements software (AR 5.20.01) and the microscope was equipped with an automated stage and automated focusing system for automated image acquisition.

### Sequencing

Sequencing was performed as described previously, with a manual first strand synthesis step and then sequencing with an altered sequencing recipe using a MiSeq sequencer (Illumina)^14,15^. The runs were single read with 40 cycles.

### Extraction of observed dwell times

Observed dwell times were extracted using the python package Papylio (https://github.com/Chirlmin-Joo-lab/papylio). First, a spatial background correction was applied using a 20-pixel median filter on the 20-frame averaged image. From the corrected averaged image, molecules were localized by finding the local maxima, discarding molecules close to the edge of the image and molecules which could not be fit with a 2D-Gaussian. Next, traces were extracted for each molecule using a Gaussian mask.

For the SPARXS-Hyb experiments, sequence identification and coupling of the single-molecule and sequencing data were also performed with Papylio. Only molecules with a five-frame rolling average intensity below a set threshold, the expected maximum intensity of a single molecule, were kept. Traces were then fit with a two-state hidden Markov model, or when no binding events were detected with a one-state Gaussian distribution. The low intensity state was classified as bound, and the high intensity state was classified as unbound. Bound (*τ_bound_*) and unbound dwell times (*τ_unbound_*) were determined from the classified traces, discarding events interrupted by the start or end of the movie, and by fitting the distribution with a single- or double-exponential decay.

### Thermodynamic and kinetic analysis

All kinetic modelling was performed using custom-written Python (version 3.12) scripts. Thermodynamic parameters for DNA duplex formation were calculated using the NUPACK Python^19^ (v4.0.2.0) with reaction conditions set to 25 °C, 0.5 M Na^+^, 0 M Mg^2+^ and a strand concentration of 10 nM. For each sequence *s*, its Watson–Crick reverse complement *s̄* was generated, and the duplex formation reaction was evaluated via partition function calculations, yielding the free energy change upon hybridization *ΔG*(*s, s̄*).

Off-target binding was quantified by evaluating each sequence’s reverse complement against all library sequences. The number of binding-competent strands was determined as:

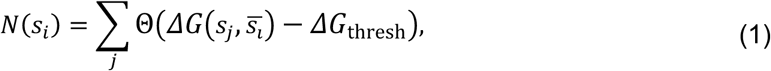

with *ΔG*_thresh_ = −14.5 k_B_T, chosen to define sequences that are thermodynamically stable within the time resolution, and Θ being the Heaviside function.

The apparent association rate was computed from *τ_unbound_* using:

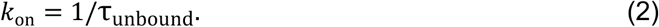

Assuming independent parallel binding pathways, the intrinsic bimolecular hybridization rate constant was obtained according to:

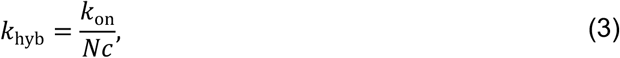

with *c* = 10 nM being the imager strand concentration.

### Cell culture

COS-7 cells (DSMZ GmbH) were cultured in Dulbecco’s modified Eagle medium (DMEM) in high glucose (Thermo Fisher) with addition of 10% fetal bovine serum (FBS; Gibco, Thermo Fisher), 1% L-glutamine (Gibco, Thermo Fisher), 1% sodium pyruvate (Gibco, Thermo Fisher), and 1% penicillin-streptomycin (Gibco, Thermo Fisher). On the day before immunostaining, cells were seeded on coverslips (#1.5 type 25 mm diameter, Carl Roth) in 6-well plates in the culture medium and incubated at 37°C with 5% CO2.

### Microtubule immunostaining

24 hours after seeding cells, α-tubulin on microtubules were immunostained with DNA-PAINT probes. Firstly, extraction buffer based on the Kapitein microtubule buffer (80 mM PIPES, 7 mM MgCl₂, 1 mM EGTA, 150 mM NaCl, 5 mM D-glucose) supplemented with 0.3% (v/v) Triton X-100 and 0.25% (w/v) glutaraldehyde was pre-warmed at 37 °C and applied to the cells for 90 seconds at room temperature. Following removal of the extraction buffer, cells were fixed for 10 minutes at room temperature with a fixation buffer containing 4% (w/v) paraformaldehyde in PBS, then washed three times for 5 minutes each with PBS. To reduce autofluorescence, cells were treated for 7 minutes with freshly dissolved 10 mM sodium borohydride in PBS and immediately rinsed with PBS afterwards before two more 10-minute washes on an orbital shaker. The fixed cells were then permeabilized with 0.25% (v/v) Triton X-100 in PBS for 7 minutes and subsequently washed three times for 5 minutes each with PBS. Later, the cells were incubated for one hour at room temperature with blocking buffer containing 2% (w/v) bovine serum albumin, 10 mM glycine, and 50 mM ammonium chloride (NH₄Cl).

Primary antibody (mouse anti-α-tubulin, T5168-.2ML, Merck) was diluted 200 times in the blocking buffer and then added to the cells for one hour at room temperature followed by three washes with the same buffer. Secondary anti-mouse nanobodies were produced and conjugated with DNA docking strand PS3 (Biomers) in house. Nanobody production and site-specific nanobody-DNA conjugation via DBCO-azide click chemistry were described in detail elsewhere^26^. Docking strand SH6 conjugated with secondary anti-mouse sdAbs were custom-made (Massive Photonics). Docking strand P1 conjugated with anti-mouse sdAbs were purchased (Massive Photonics). All secondary nanobodies or antibodies were diluted with the blocking buffer (SH6 at 0.08uM, PS3 at 0.04 µM, and P1 at 0.1 µM) and applied on cells on respective coverslips during one-hour incubation, followed by three 5-minute washes with the blocking buffer.

Cells were post-fixated with 2% PFA after immunostaining and stored in PBS at 4 degrees. During imaging, coverslips were mounted with imaging buffer containing corresponding imager strands diluted at various concentrations in 500 mM NaCl solution in PBS. An overview of the DNA sequences, concentrations and imaging parameters can be found in Table S2.

### Microscopy setup for cell experiments

A custom-built TIRF microscope was used for DNA-PAINT imaging in cells^27^. Excitation from a 561 nm laser (MGL-FN-561, CNI Laser) is delivered via a multimode optical fibre (NA 0.22, square core profile of size 70 μm by 70 μm, customized, Ceram Optec) to the microscope. The laser beam is collimated by a 30 mm achromatic lens (AC254−030-A, Thorlabs) after the fibre exit and focused by a 150 mm achromatic lens (147−643, Edmund Optics) onto the back focal plane of an oil-immersion objective (NA 1.5, 60×, UPLAPO60XOHR, Olympus). A one-dimensional motorized stage (KMTS25E/M, Thorlabs) is incorporated to translate the beam position away from the optical axis for TIRF illumination. Homogeneous illumination is facilitated by a vibration motor (5 mm Vibration Motor, 304−111, Precision Microdrives) which agitates the optical fibre through a 3D printed mount. Sample positioning is achieved via a three-dimensional stick−slip piezo stage (assembled by three identical linear stages, CLS5252, Smaract). Both the sample stage and the objective are fixed on the customized MiCube microscope body^28^. Fluorescence is decoupled from the excitation beam using a quad-band dichroic mirror (zt405/488/561/ 640rpc, Chroma) and further filtered by a notch filter (ZET405/488/ 561/640mv2, Chroma). A 180 mm tube lens (TTL80-A, Thorlabs) followed by two 300 mm lenses (G322336322, Qioptiq) in 4f configuration focused the image onto an sCMOS camera (BSI Express, Photometrics). A bandpass emission filter (ET595/50m, Chroma) was inserted for fluorescence clean-up of Cy3(B).

### DNA-PAINT image analysis

Image stacks were acquired using μManager 2.0 software. DNA-PAINT images were reconstructed using Picasso software^29^ (v0.9.5). Drift correction was done using the redundant cross-correlation (RCC) algorithm^30^ integrated in Picasso. The localization precision was estimated by Picasso’s integrated nearest neighbour based analysis (NeNA) algorithm^31^. To analyse localization density (Fig. 4b), five circular regions of interest (ROIs) with a diameter of 5.4 μm covering similar filament density were selected per DNA-PAINT image. The average number of localizations per image frame was extracted from every ROI and plotted using Python.

## Author contributions

CB developed, performed and analysed the SPARXS-Hyb experiments under supervision of CJ. AS performed the serial control experiments. RH performed and analysed the cellular imaging under supervision of KSG. PI developed and performed the thermodynamic and kinetic analysis under supervision of RS. CB, RH, PI, KSG, and CJ wrote the manuscript.

## Data and code availability

The data supporting the findings of this study are available in the article and in the supplementary information. Additional data and the custom analysis code are available from the corresponding author upon request.

## Acknowledgements

We thank Mike Filius for fruitful discussions, Ivo Severins for help with the software, and Martin Depken and Hidde Offerhaus for their advice on the data analysis. CB received seed funding from the Kavli Synergy program of the Kavli Institute for Nanoscience Delft. The work was also made possible with the support of a postdoc fellowship from the German Academic Exchange Service (DAAD) to PI. This work was supported by the HORIZON-MSCADN-2022 (DYNAMO, 101072818); by the Basic Research Laboratory Program (grant no. RS-2026-25485403 to CJ); by the Ministry of Science and ICT (Bio&Medical Technology Development Program of the National Research Foundation, RS-2025-02217909 to CJ); by the TU Delft Department of Bionanoscience (startup fund to KSG), by the TU Delft Open Research Hardware Stimulation Fund to RH; by the Netherlands Organization for Scientific Research (NWO) (Vidi grant, VI.Vidi.243.206 to KSG); by an ERC grant (QScope, 101165129 to KSG). Views and opinions expressed are however those of the author(s) only and do not necessarily reflect those of the European Union or the European Research Council Executive Agency. Neither the European Union nor the granting authority can be held responsible for them.

## Supplementary information

**Fig. S1.**
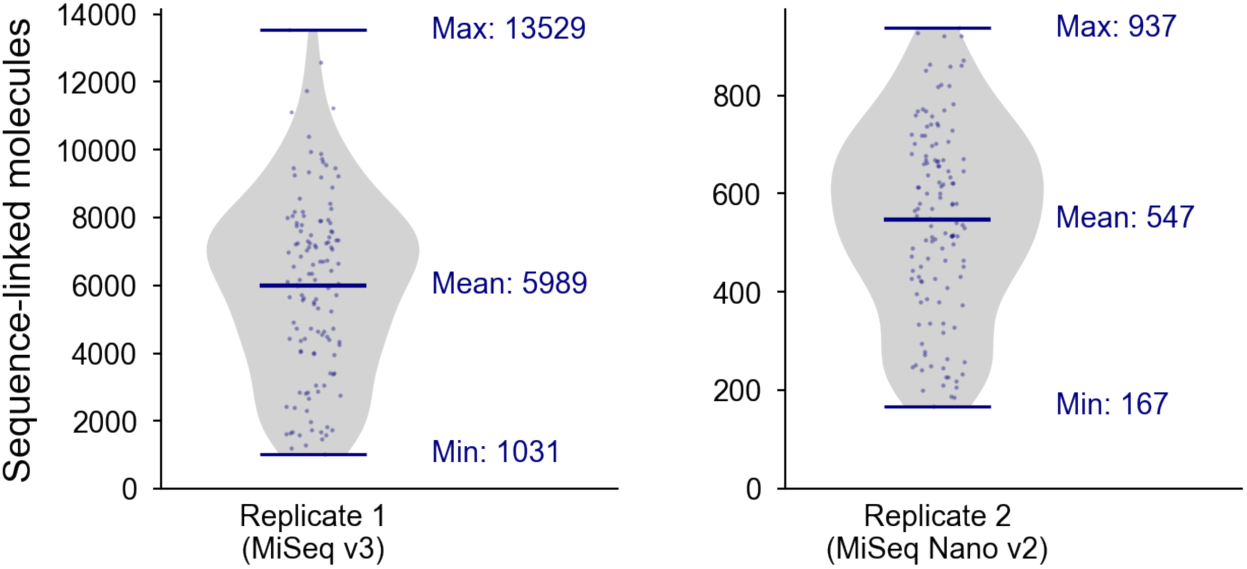
Violin plots of sequence-linked molecules for each SPARXS-Hyb replicate. Replicate 1 and 2 were performed using a v3 and v2 Nano MiSeq sequencing flow cell, respectively. One sequence (GAGGAGG) was spiked in and was therefore much more abundant (20,555 molecules) than the others and excluded from the plot.

**Fig. S2.**
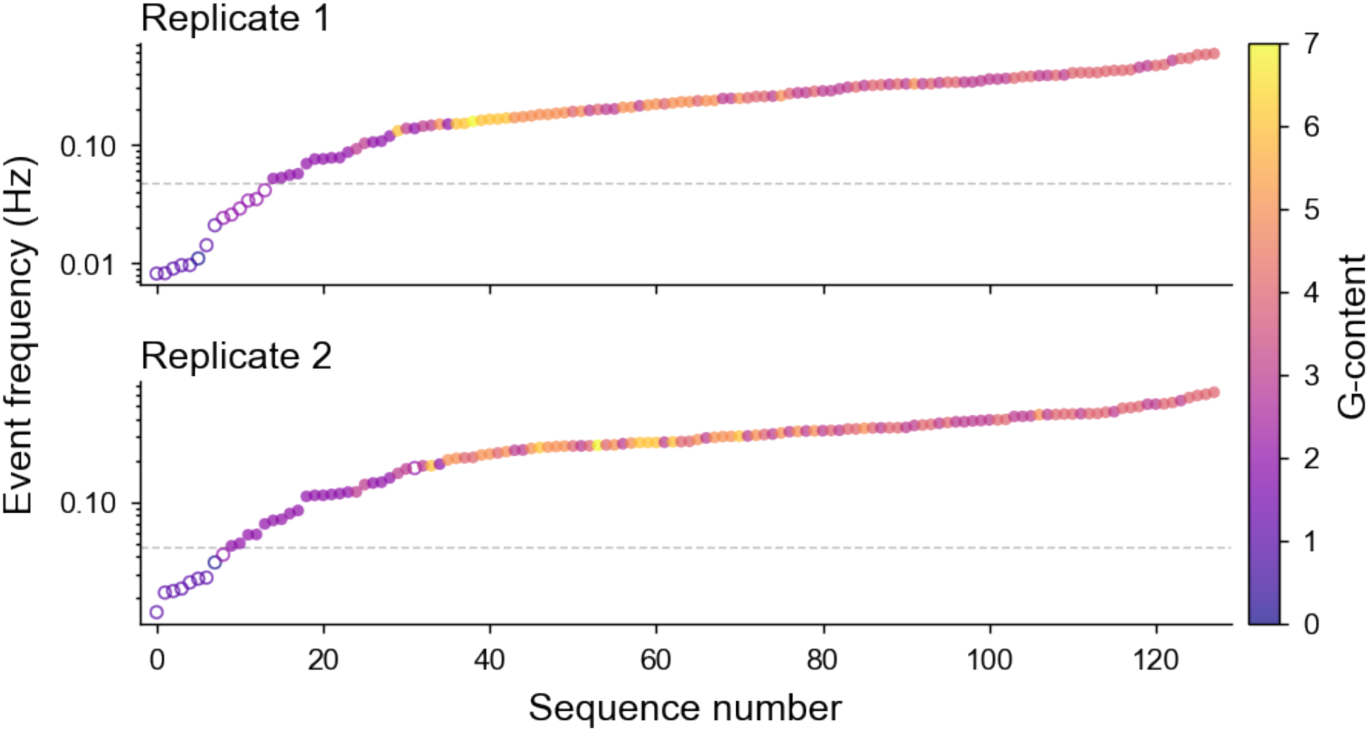
Sequences ranked by event frequency. Sequences with an event frequency below 0.05 Hz (the dashed line) were excluded from further analysis. The colour of the data points indicates the number of guanine nucleotides in the docking sequence. Replicate 1 and 2 were performed using a v3 and v2 Nano MiSeq sequencing flow cell, respectively. For each replicate all 128 library sequences are shown.

**Fig. S3.**
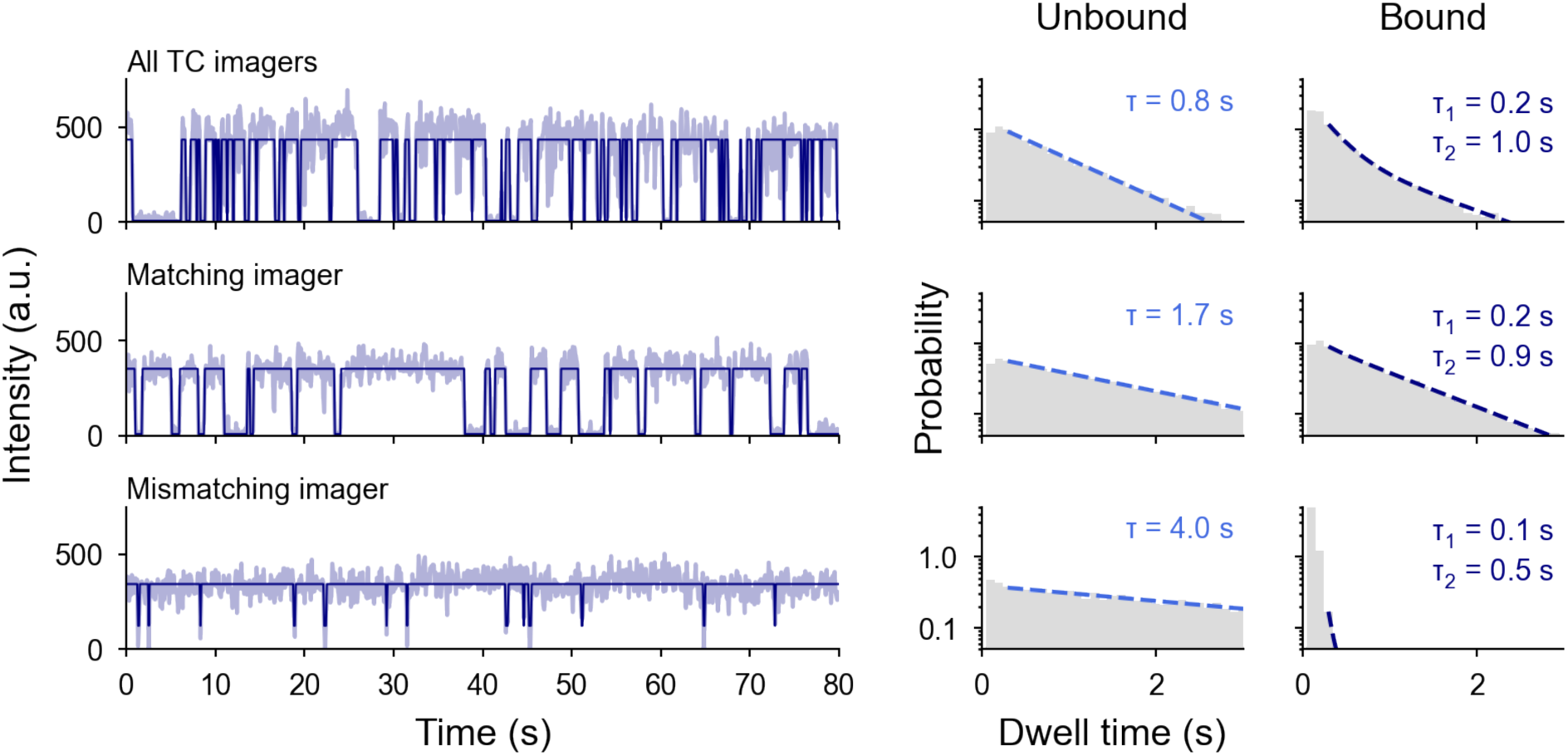
Examples of matching and mismatching imager traces and distributions. **a,** Representative fluorescence time traces (light) with hidden Markov model classification (dark) of a single docking sequence which was immobilized on a custom flow cell. From top to bottom, the solution contained: 10 nM of each TC-imager (i.e. the imager library at 1.3 µM), 10 nM of the matching imager (SH1), 10 nM of a single mismatching imager (Imager for AAGGGGA). **b,** Probability density distributions of the bound and unbound dwell times for, from top to bottom, the combination of all TC-imagers (n = 18,903 and 18,787), the matching imager (n = 44,397 and 45,281), and a mismatching imager (n = 15,060 and 16,709).

**Fig. S4.**
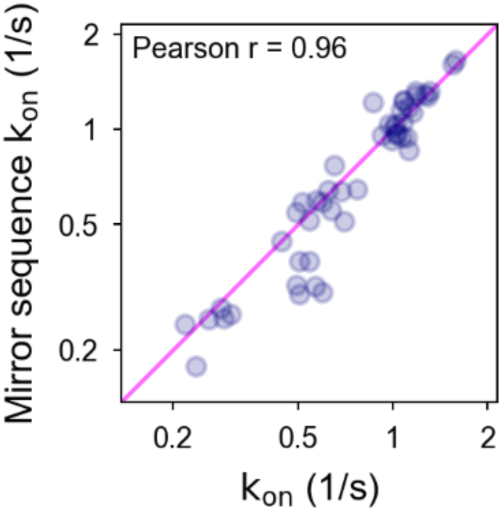
Correlation of observed association rates for mirror sequences. Mirror sequences are pairs of sequences where the 5’-3’ direction of one sequence equals the 3’-5’ direction of the other. Sequences with an event frequency below 0.05 Hz were excluded, leaving 48 mirror pairs.

**Fig. S5.**
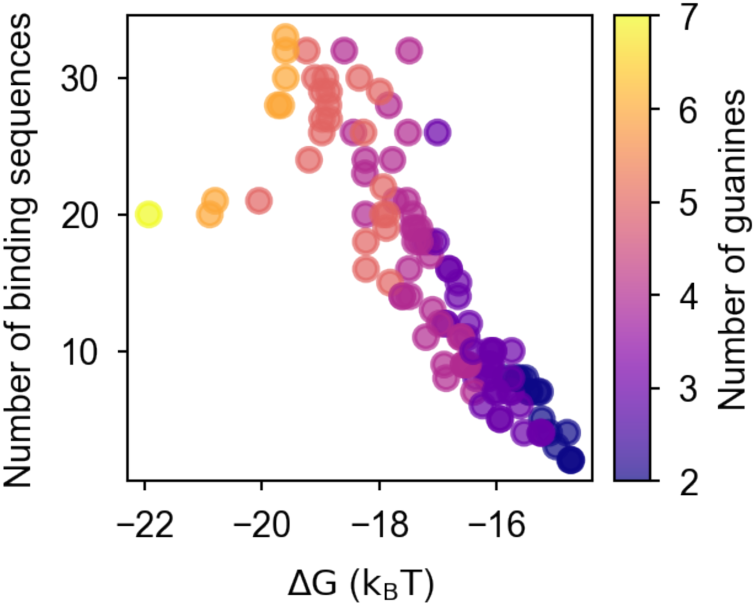
Number of imager sequences that can bind each docking sequence. The free-energy change of hybridizing every imager sequence to each docking sequence was calculated using NUPACK^19^. Imager sequences were counted as potential binding sequences when the free-energy change was below −14.5 k_B_T. The colour of the data points indicates the number of guanine nucleotides in the docking sequence. All 128 library sequences are shown.

**Fig. S6.**
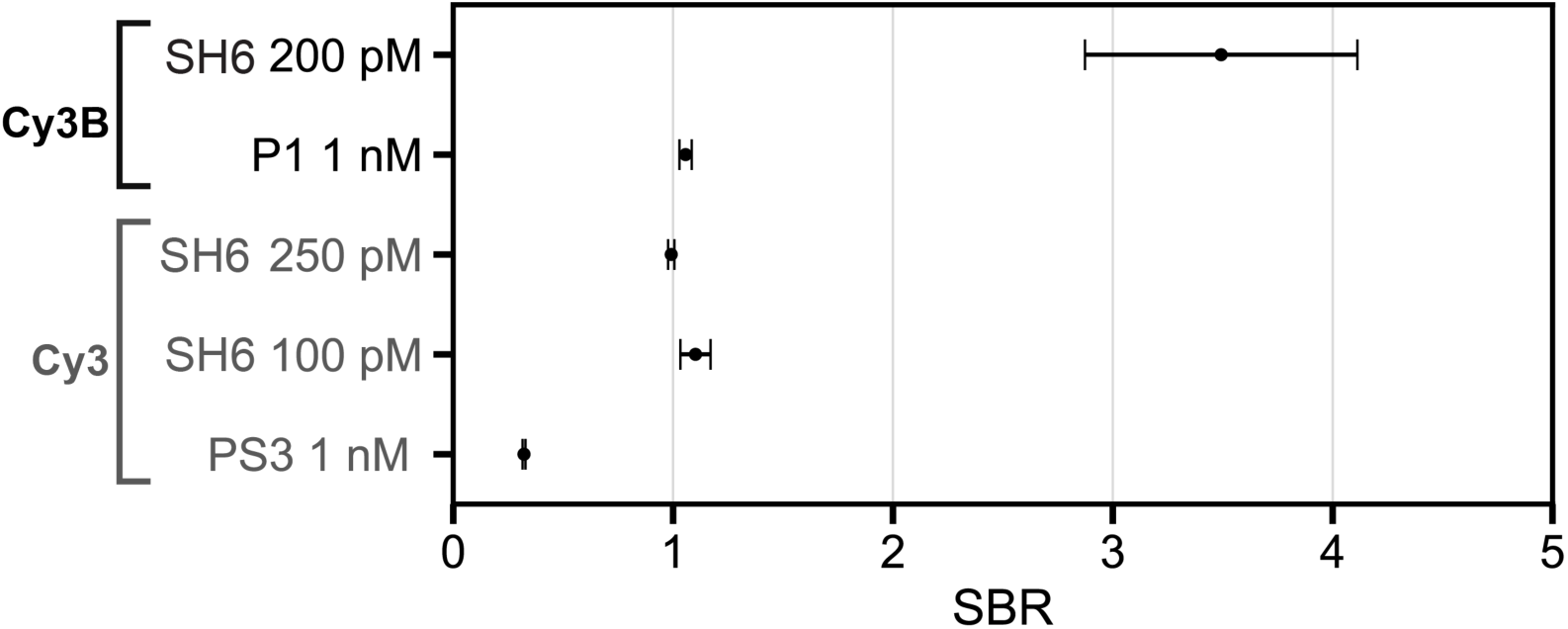
Average signal-to-background ratio (SBR) of cell images compared between. 1 nM of PS3, 100 pM of SH6, and 250 pM of SH6, which were imaged using Cy3 fluorophore; 1 nM of P1, and 200 pM of SH6, which were imaged using brighter Cy3B fluorophore. SBR is calculated for each localization by the ratio of the peak intensity and the background in number of photons, whereas the peak intensity is defined by the one-pixel area integral over the point spread function described by the Gaussian fit parameters, and the background is the offset value per pixel ^32^. Error bar denotes the standard deviation of SBR of localizations from five regions of interest in each DNA-PAINT image.

**Table S2.** Summary of sample and image parameters.

| Figure | Imager type and concentration | Laser intensity (kW/cm<sup>2</sup>) | Camera exposure time (ms) | Number of frames | NeNA localization precision (nm) |
| --- | --- | --- | --- | --- | --- |
| Fig.4b(i) | PS3-Cy3, 1 nM | 0.46 | 50 | 10000 | 14.6 |
| Fig.4b(ii) | SH6-Cy3, 100 pM | 0.30 | 50 | 10000 | 10.9 |
| Fig.4b(iii) | SH6-Cy3, 250 pM | 0.39 | 50 | 10000 | 11.7 |
| Fig.4a | SH6-Cy3B, 200 pM | 0.39 | 50 | 10000 | 7.82 |
| Fig. S6 | P1-Cy3B, 1 nM | 0.39 | 100 | 10000 | 9.43 |

**Table S3.**
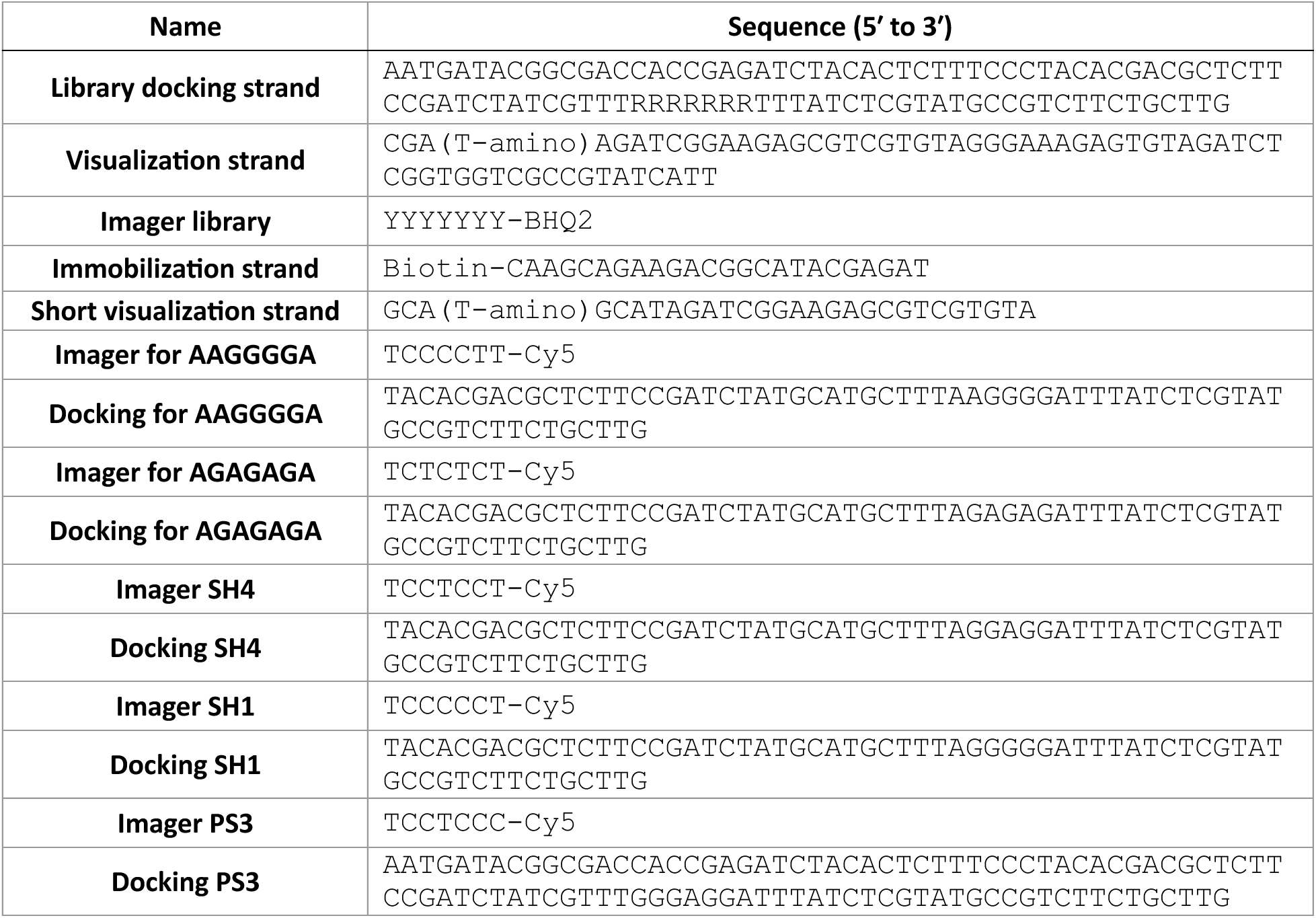
Sequences of DNA oligonucleotides used for SPARXS-Hyb and controls.

## References

1. Seeman, N. C. & Sleiman, H. F. DNA nanotechnology. Nat. Rev. Mater. 3, 17068 (2017).

2. Zhan, P. et al. Recent Advances in DNA Origami-Engineered Nanomaterials and Applications. Chem. Rev. 123, 3976–4050 (2023).

3. Yang, S. et al. DNA as a universal chemical substrate for computing and data storage. Nat. Rev. Chem. 8, 179–194 (2024).

4. Jungmann, R. et al. Single-Molecule Kinetics and Super-Resolution Microscopy by Fluorescence Imaging of Transient Binding on DNA Origami. Nano Lett. 10, 4756–4761 (2010).

5. Strauss, S. & Jungmann, R. Up to 100-fold speed-up and multiplexing in optimized DNA-PAINT. Nat. Methods 17, 789–791 (2020).

6. Schueder, F. et al. An order of magnitude faster DNA-PAINT imaging by optimized sequence design and buffer conditions. Nat. Methods 16, 1101–1104 (2019).

7. Banerjee, A., Anand, M., Srivastava, M., Vidwath, V. S. & Ganji, M. High-speed multiplexed DNA-PAINT imaging of nuclear organization using an expanded sequence repertoire. Nat. Commun. 17, 3655 (2026).

8. Helmerich, D. A. et al. Impact of Docking Strand Design on Spatial Resolution in DNA-Points Accumulation for Imaging in Nanoscale Topography. ChemPhysChem 27, e202500803 (2026).

9. SantaLucia, J. & Hicks, D. The Thermodynamics of DNA Structural Motifs. Annu. Rev. Biophys. Biomol. Struct. 33, 415–440 (2004).

10. Zadeh, J. N. et al. NUPACK: Analysis and design of nucleic acid systems. J. Comput. Chem. 32, 170–173 (2011).

11. Hata, H., Kitajima, T. & Suyama, A. Influence of thermodynamically unfavorable secondary structures on DNA hybridization kinetics. Nucleic Acids Res. 46, 782–791 (2018).

12. Zhang, J. X. et al. Predicting DNA hybridization kinetics from sequence. Nat. Chem. 10, 91–98 (2018).

13. Hertel, S. et al. The stability and number of nucleating interactions determine DNA hybridization rates in the absence of secondary structure. Nucleic Acids Res. 50, 7829–7841 (2022).

14. Severins, I. et al. Single-molecule structural and kinetic studies across sequence space. Science 385, 898–904 (2024).

15. Bastiaanssen, C., Severins, I., Van Noort, J. & Joo, C. Single-molecule parallel analysis for rapid exploration of sequence space. Nat. Protoc. https://doi.org/10.1038/s41596-025-01196-y (2025) doi:10.1038/s41596-025-01196-y.

16. Aguirre Rivera, J., et al. Massively parallel analysis of single-molecule dynamics on next-generation sequencing chips. Science 385, 892–898 (2024).

17. Panfilov, M. et al. Multiplexed single-molecule characterization at the library scale. Nat. Protoc. 21, 749–774 (2026).

18. Gao, Y. Secondary structure effects on DNA hybridization kinetics: a solution versus surface comparison. Nucleic Acids Res. 34, 3370–3377 (2006).

19. Fornace, M. E. et al. NUPACK: Analysis and Design of Nucleic Acid Structures, Devices, and Systems. Preprint at 10.26434/chemrxiv-2022-xv98l (2022).

20. Li, J. et al. Exploring the speed limit of toehold exchange with a cartwheeling DNA acrobat. Nat. Nanotechnol. 13, 723–729 (2018).

21. Gowri, G., Sheng, K. & Yin, P. Scalable design of orthogonal DNA barcode libraries. Nat. Comput. Sci. 4, 423–428 (2024).

22. Dertinger, T., Colyer, R., Iyer, G., Weiss, S. & Enderlein, J. Fast, background-free, 3D super-resolution optical fluctuation imaging (SOFI). Proc. Natl. Acad. Sci. 106, 22287–22292 (2009).

23. Glogger, M., Spahn, C., Enderlein, J. & Heilemann, M. Multi-Color, Bleaching-Resistant Super-Resolution Optical Fluctuation Imaging with Oligonucleotide-Based Exchangeable Fluorophores. Angew. Chem. Int. Ed. 60, 6310–6313 (2021).

24. Wade, O. K. et al. 124-Color Super-resolution Imaging by Engineering DNA-PAINT Blinking Kinetics. Nano Lett. 19, 2641–2646 (2019).

25. Chandradoss, S. D. et al. Surface passivation for single-molecule protein studies. J. Vis. Exp. https://doi.org/10.3791/50549 (2014) doi:10.3791/50549.

26. De Zwaan, K. et al. High-Throughput Single-Molecule Microscopy with Adaptable Spatial Resolution Using Exchangeable Oligonucleotide Labels. ACS Nano 19, 13149–13159 (2025).

27. de Zwaan, K. et al. High-Throughput Single-Molecule Microscopy with Adaptable Spatial Resolution Using Exchangeable Oligonucleotide Labels. ACS Nano 19, 13149–13159 (2025).

28. Martens, K. J. A. et al. Visualisation of dCas9 target search in vivo using an open-microscopy framework. Nat. Commun. 10, 3552 (2019).

29. Schnitzbauer, J., Strauss, M. T., Schlichthaerle, T., Schueder, F. & Jungmann, R. Super-resolution microscopy with DNA-PAINT. Nat. Protoc. 12, 1198–1228 (2017).

30. Wang, Y. et al. Localization events-based sample drift correction for localization microscopy with redundant cross-correlation algorithm. Opt. Express 22, 15982 (2014).

31. Endesfelder, U., Malkusch, S., Fricke, F. & Heilemann, M. A simple method to estimate the average localization precision of a single-molecule localization microscopy experiment. Histochem. Cell Biol. 141, 629–638 (2014).

32. Steen, P. R. et al. The DNA-PAINT palette: a comprehensive performance analysis of fluorescent dyes. Nat. Methods 21, 1755–1762 (2024).

